# ESCO: single cell expression simulation incorporating gene co-expression

**DOI:** 10.1101/2020.10.20.347211

**Authors:** Jinjin Tian, Jiebiao Wang, Kathryn Roeder

## Abstract

**Motivation:** Gene-gene co-expression networks (GCN) are of biological interest for the useful information they provide for understanding gene-gene interactions. The advent of single cell RNA-sequencing allows us to examine more subtle gene co-expression occurring within a cell type. Many imputation and denoising methods have been developed to deal with the technical challenges observed in single cell data; meanwhile, several simulators have been developed for benchmarking and assessing these methods. Most of these simulators, however, either do not incorporate gene co-expression or generate co-expression in an inconvenient manner.

**Results:** Therefore, with the focus on gene co-expression, we propose a new simulator, ESCO, which adopts the idea of the copula to impose gene co-expression, while preserving the highlights of available simulators, which perform well for simulation of gene expression marginally. Using ESCO, we assess the performance of imputation methods on GCN recovery and find that imputation generally helps GCN recovery when the data are not too sparse, and the ensemble imputation method works best among leading methods. In contrast, imputation fails to help in the presence of an excessive fraction of zero counts, where simple data aggregating methods are a better choice. These findings are further verified with mouse and human brain cell data.

**Availability:** The ESCO implementation is available as R package `SplatterESCO` (https://github.com/JINJINT/SplatterESCO).

## 1 Introduction

A synchronization between gene expression leads to gene co-expression. Cell heterogeneity, due to cell type or cell cycle, can generate correlations between genes that are highly expressed in similar cells. Alternatively, any form of gene cooperation within a cell type, such as gene co-regulation, also results in co-expression. To differentiate these two settings, we refer them as the gene co-expression across heterogeneous cell groups and gene co-expression within homogeneous cell groups respectively, throughout this article. Understanding gene co-expression in the former setting helps with cell-type identification, and in the latter setting, it helps detect gene regulation relationships and can further provide insights into genetic disorders (Pang *et al.*, 2020; Polioudakis *et al.*, 2019; Parikshak *et al.*, 2013; Willsey *et al.*, 2013).

Single-cell RNA sequencing (scRNA-seq), a recent breakthrough technology that paves the way for measuring transcription at single cell resolution to study precise biological functions, allows us to target gene co-expression within homogeneous cell groups for the first time. Indeed, early statistical models argued that genes within homogeneous cell groups were independent (Quinn *et al.*, 2018). However, they overlooked the investigations from the biological end, which reveal that correlation arises due to the stochastic nature of gene expression and gene regulation dynamics (Raj *et al.*, 2006).

scRNA-seq data present many challenges for co-expression analysis, due to the sparsity of counts, which include many zeros, mainly arising from low capture and sequencing efficiency in the data collecting process. Sparsity occurs in both a gene- and a cell-specific manner and is observed to have the greatest impact on genes that have low expression. An ever-growing literature attempts to address these challenges using imputation and other denoising methods (Chen *et al.*, 2020; Gong *et al.*, 2018; Huang *et al.*, 2018; Li and Li, 2018; Van Dijk *et al.*, 2018; Eraslan *et al.*, 2019; Linderman *et al.*, 2018). To systemically benchmark these methods, we require realistic simulation tools to construct a ground truth for scRNA-seq data with realistic technical noise; however, currently there is a paucity of methods for this purpose.

Numerous scRNA-seq simulators using both non-parametric and parametric approaches have been proposed during recent years, e.g., Splat (Zappia *et al.*, 2017), SymSim (Zhang *et al.*, 2019a), PROSSTT (Papadopoulos *et al.*, 2019), and SERGIO (Dibaeinia and Sinha, 2020). Each of those methods focuses on producing realistic marginal behavior of gene expression, and successfully modeling these features, as well as capturing cell type heterogeneity. But, those simulators either ignore gene co-expression, or they generate it in a way that is hard to benchmark. Real data clearly display gene co-expression within homogeneous cell groups (**Fig. 1A**) and gene co-expression across heterogeneous cell groups (**Fig. 1B**). By contrast, almost all gene pairs show no correlation for simulated data generated using Splat, even without the challenge of added technical noise (**Fig. 1C**). While the data simulated by SymSim may show a modest level of gene co-expression (**Fig. 1D** left panel), that correlation arises from the cell type confounding^1^, rather than true gene-gene interaction (**Fig. 1D** right panel).

**Fig. 1.**
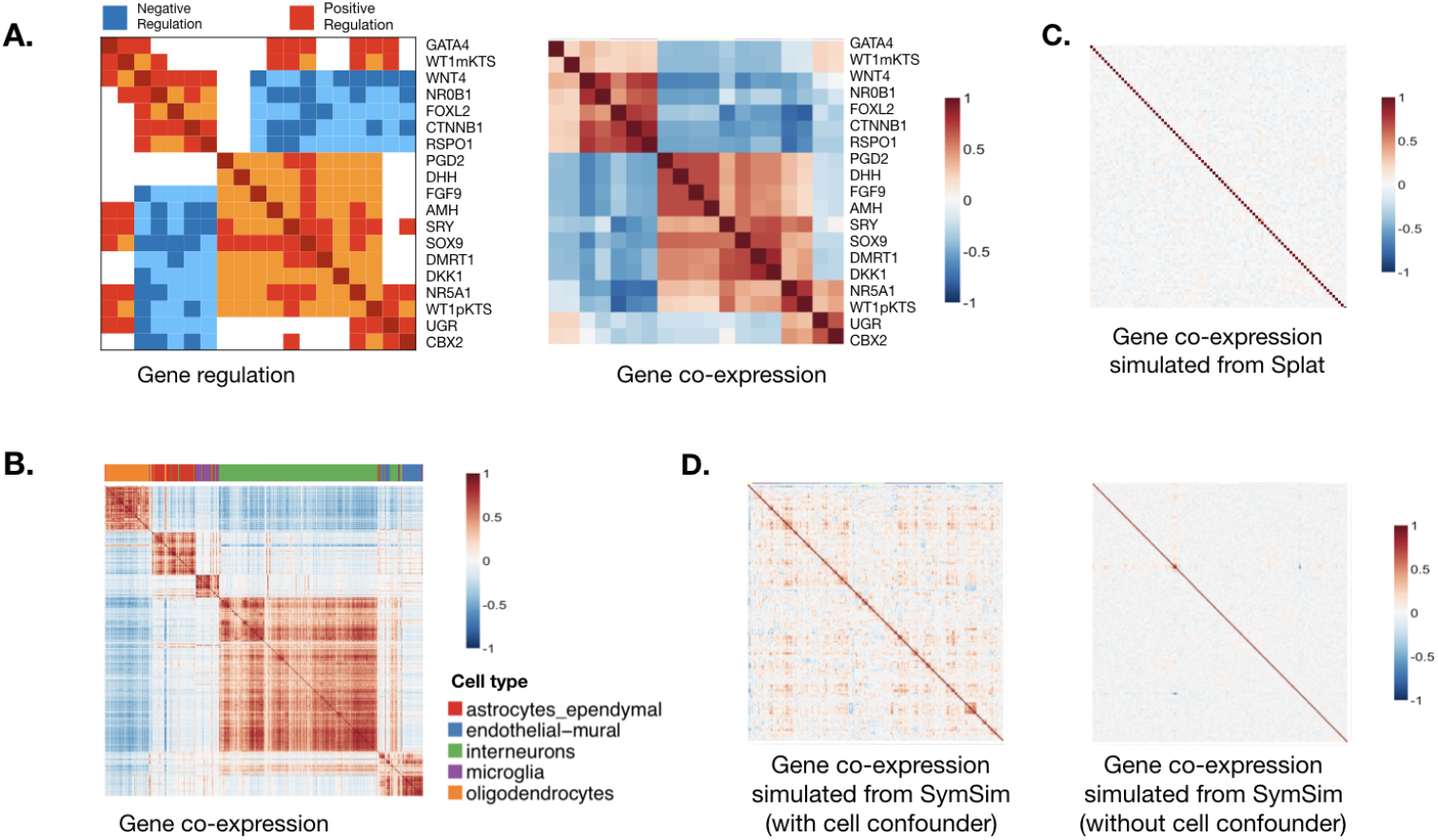
Gene co-expression is informative, but we lack satisfactory methods to simulate it for scRNA-seq data. **A.** Connection between gene regulation and gene co-expression. The left panel shows the regulation relationship between the 19 genes in Gonadal Sex Determination (Ríos et al., 2015), while the right panel shows Pearson’s correlation matrix for these 19 genes with inferred expression (Pratapa et al., 2020). **B.** Connection between gene co-expression and cell group clusters. The correlation matrix of the 500 most significant marker genes of the five major cell types from the Zeisel data (Zeisel et al., 2015) with corresponding gene types marked with a color bar on top, clustered using hierarchical clustering. **C.** The correlation matrix for 200 simulated genes from Splat (Zappia et al., 2017), without zero-inflation. **D.** The correlation matrix for 200 simulated genes from SymSim (Zappia et al., 2017), without zero-inflation. The left and right panels show results with and without the cell confounding effect, respectively.

Here we propose a new simulation tool, **E**nsemble **S**ingle-cell expression simulator incorporating gene **CO**-expression, ESCO, which is constructed as an ensemble of the best features among current simulators to preserve the marginal performance, while allowing easily incorporating co-expression structure among genes using a copula. Particularly, ESCO allows realistic simulation of a homogeneous cell group, heterogeneous cell groups, as well as complex cell group relationships such as tree and trajectory structure, together with a flexible input of co-expression. As for technical noise, ESCO integrates the parametric and non-parametric approaches in current literature and gives the user flexibility to choose. In order to mimic a specific real data set, ESCO can estimate all the hyperparameters in a feasible way for both a homogeneous cell group or heterogeneous cell groups. ESCO is implemented in the R package `SplatterESCO`, which is built upon the R package `Splatter` (Zappia *et al.*, 2017), in order to provide a unified software framework.

## 2 Methods

### 2.1 Models

Despite their differences, current simulation approaches arguably follow a general flowchart (**Fig. 2**). For example, Splat (Zappia *et al.*, 2017) simulates scRNA-seq data using a hierarchical model in which the gamma-Poisson distribution imposes a mean and variance trend; SymSim (Zhang *et al.*, 2019a) is based on a similar hierarchical model with gene kinetics guiding the hyperparameter selection, a non-parametric approach to introduce more realistic noise, and a focus on tree-structured heterogeneity; PROSSTT (Papadopoulos *et al.*, 2019) aims to simulate realistic cell trajectories using a model based on Brownian motion; SERGIO (Dibaeinia and Sinha, 2020) starts from the gene regulation relationship and solves a series of stochastic differential equations given by gene kinetics to impose those regulations. The more complex non-parametric modeling tends to fit data better than parametric modeling, given that the aim is to mimic data for which the model has already been trained. However, this approach is not practical for producing simulated data similar to a new data set. For example, the non-parametric methods like SymSim and SERGIO use grid search over a large number of tuning parameters. By contrast, the parametric Splat approach can be tuned to data by fitting a one-step statistical regression model. ESCO also follows the general flowchart in **Fig. 2**, but it aims to incorporate the best features from the existing methods. **Fig. 3** demonstrates the superiority of ESCO, as it allows simulation of scRNA-seq data with various cell heterogeneity and customized gene co-expression patterns. In this section, we elaborate on the specific simulation models that ESCO adopts, following the framework outlined in **Fig. 2**.

**Fig. 2.**
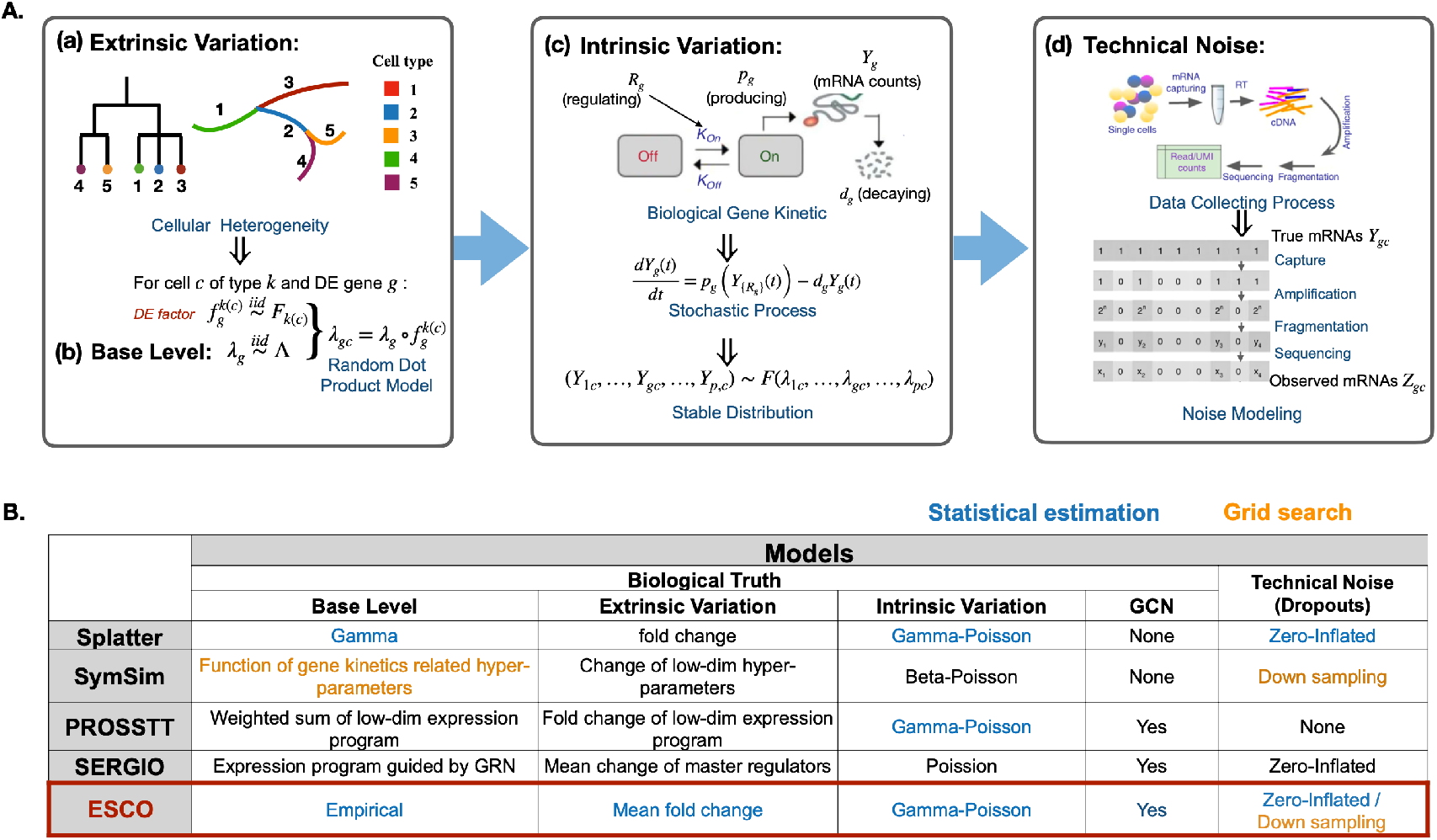
Summary of simulators for scRNA-seq data. **A.** The general modeling flowchart of commonly used simulators. Simulators often start with **(a)** extrinsic variation that arose from cell heterogeneity in the biological sense, and import this model to **(b)** the base expression mean generated for each gene, to formalize the heterogeneous expression means for a gene in a cell of a particular cell type. Then, those means are used to generate the expression level, i.e., mRNA counts, by modeling the **(c)** intrinsic variation, i.e., the stochasticity of gene expression in a cell with a defined base rate of expression. This process is often modeled by the gene kinetic model in biochemistry, which could be stated as a stochastic process in statistical terms. The stable distribution of this stochastic process can usually be approximated by distributions like negative binomial / Poisson / beta Poisson. Finally, some simulators allow the generation of technical noise **(d)** separately, by adding noise, step by step, to the true counts, to mimic the data collection process (the cartoon display is from Zhang et al. (2019a)). Usually, this stepwise process is approximated by the zero-inflation model, where the true counts are set to zero with probability related to expression level. **B.** Summary of the current state of simulators following the general modeling flowchart described above, with blue and orange text color indicating whether they use statistical estimation or grid search when fitting the simulator to a real data set. The objective of ESCO is to create an ensemble of the best features among current simulators in each step, while allowing easily imposing co-expression structure among genes via a copula.

**Fig. 3.**
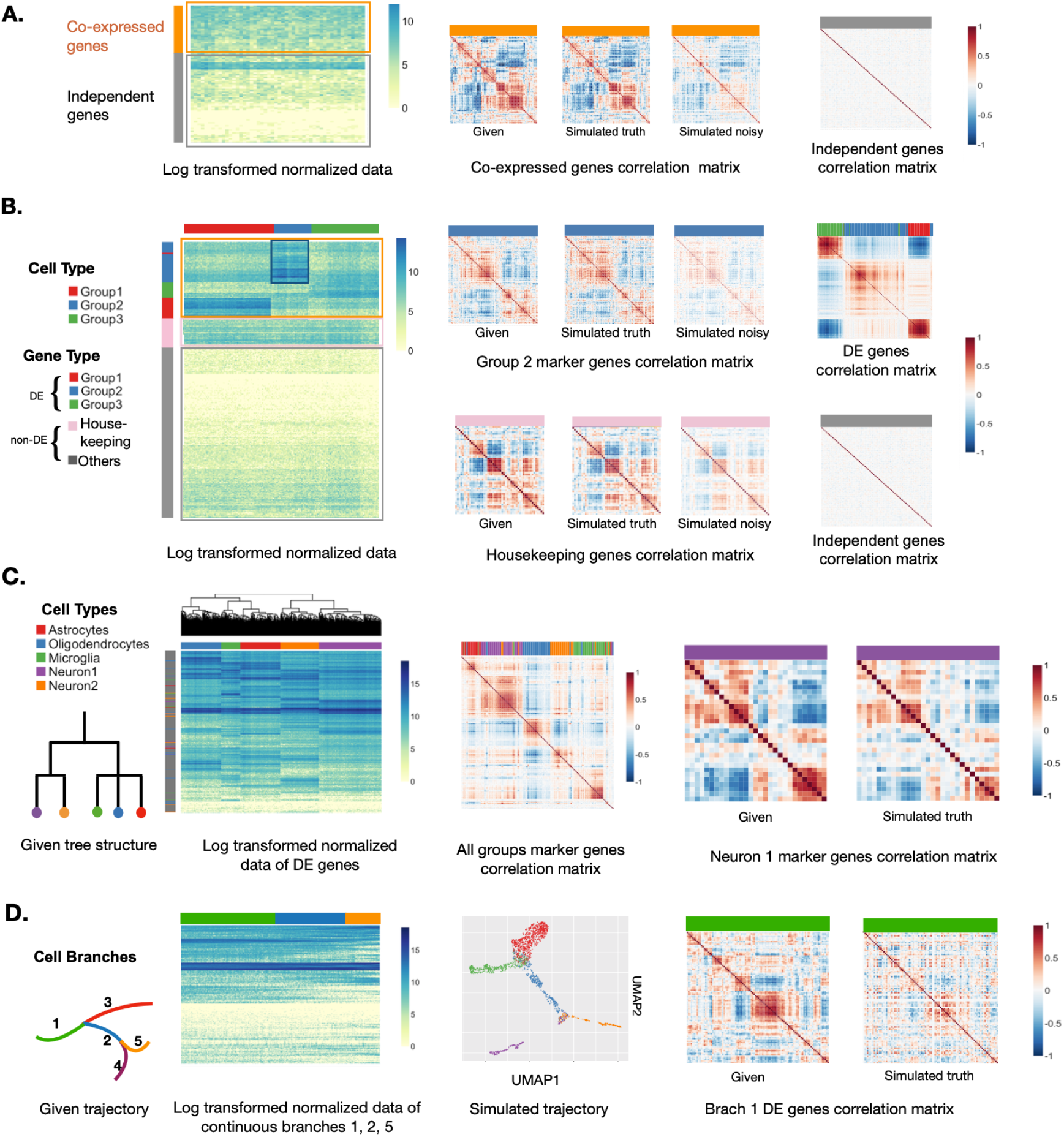
ESCO can simulate scRNA-seq data of various cell heterogeneity and gene co-expression. **A.** The simulation results for one homogeneous cell group consisting of 200 cells and 500 genes. The first panel displays the heatmap of log2 transformed normalized simulated expression data, where rows represent genes and columns represent cells; 30% of genes are chosen to be co-expressed genes, and the rest are independent genes. The following displays depict, in order, the given correlation structure for co-expressed genes, the simulated correlation structure among those co-expressed genes without noise, and that with technical noise, and the simulated correlation structure for independent genes. **B.** The simulation results for three discrete heterogeneous cell groups consisting of 500 cells and 1000 genes. 30% of the genes are chosen to be cell-type DE genes and presumably co-expressed, among which each marks one cell type. Another 10% of genes are chosen to be housekeeping genes, and also presumably co-expressed. The rest are independent non-DE genes. The first display shows the heatmap of log2 transformed normalized simulated data, where different gene types (rows) and cell types (columns) are marked with color bars on the margin. The following displays depict, in order in each row, the given correlation structure for both marker genes of Group2 and co-expressed housekeeping genes, the simulated correlation structure among those co-expressed genes without noise, and that with technical noise; and, at the end of each row the simulated correlation structure among all DE genes across all cells, and that among all independent genes across all cells, with corresponding gene types marked with a color bar on top. **C.** The simulation results for five heterogeneous cell groups that follow a tree structure given in the first panel. We simulate 1000 cells and 2000 genes: 30% of genes are chosen to be DE genes and presumably co-expressed, among which 5% are markers; the rest are independent non-DE genes. The second panel shows the heatmap of log2 transformed normalized simulated data. Different cell types are marked with color bars on the column margin, together with the hierarchical clustering of cells. The following displays depict, in order, the resulting correlation structure among all marker genes across all cells, with corresponding gene types marked with a color bar on top; the given correlation structure for co-expressed marker genes of Neuron1 cells, and the resulting correlation structure among those co-expressed genes. **D.** The simulation results for five heterogeneous cell groups that follow a smooth cell trajectory structure given in the first panel. There are 1000 cells and 2000 genes; 30% of genes are chosen to be DE genes and presumably co-expressed, and the rest are independent non-DE genes. The following displays depict, in order, the heatmap of log2 transformed normalized simulated data for all DE genes in one continuous path (i.e., branches 1 → 2 → 5), with branch ID marked with a color bar on top; the UMAP for the first two dimensions of the simulated data; the given correlation structure for all DE genes across one branch (branch 1), and the resulting correlation structure simulated of those genes.

#### Base expression level

We simulate base expression level in an empirical way that allows inputting any density function, either non-parametric or parametric. Particularly, we denote the base expression level for gene *g* as *λ*_*g*_, and we let

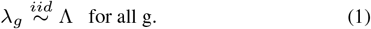

#### Extrinsic variation

The heterogeneity of cell groups is driven by the differential expressed (DE) behavior among certain gene sets across groups. Therefore we implement the cell group heterogeneity, i.e., the extrinsic variation, via modeling the behavior of DE genes. We use the random dot product model to introduce this heterogeneity by imposing a DE factor generated separately on the otherwise homogeneous gene expression means. Particularly, we generate the different cell group structures we want, via modeling the DE factor in each of the following ways.

##### 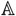. Discrete cell groups

In order to generate clear and distinguishable cell groups, we randomly split the set of DE genes into subsets, each is identified as marker genes for a cell group. Then we simulate the DE factor for each marker gene set as a LogNormal random variable with different mean and variance indexed by group identity.

Particularly, denote the set of DE genes as *G*^DE^, and the marker gene set 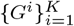 for *k* cell groups such that *G*^1^ ∪ *G*^2^ · · · ∪ *G*^*k*^ ∪ … *G*^*K*^ = *G*^DE^, we let the DE factor for each DE gene *g* in cell group *k* be

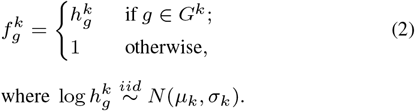

##### 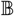. Tree-structured cell groups

We utilize the idea in SymSim (Zhang *et al.*, 2019a), which makes the DE factor of similar cell groups more related to each other. Particularly, we generate the DE factor from a multivariate normal distribution, where the covariance matrix is given by the tree structure of the data. Additionally, in order to assure the identifiability of different cell groups, we introduce extra heterogeneity via strengthening the DE factor for a small proportion of DE genes, which are identified as marker genes in this setting (different from those in the discrete cell group setting).

Specifically, given the similarity between cell groups by a *K* × *K* correlation matrix Σ generated from the tree structure, and a set of DE genes *G*^DE^, we firstly select a small proportion of *G*^DE^ and split them into the marker genes for each group *G*^1^, *G*^2^, …, *G*^*K*^. We let the DE factor for each DE gene *g* in cell group *k* be

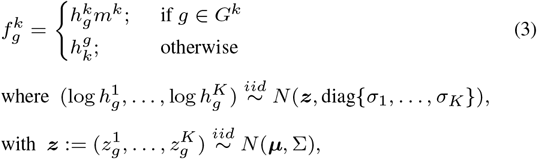

and *m*^*k*^ > 1 is a scalar parameter controlling the level of the additional heterogeneity for each group.

##### 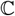. Continuous cell trajectories

We utilize the idea in PROSSTT (Papadopoulos *et al.*, 2019), which uses Brownian motion to generate the DE factors, so that the smooth cell heterogeneity can be generated. Particularly, for each gene in the DE gene set *G*^DE^, we simulate the DE factor at each step *t* in branch *b* with length *T*_*b*_ as

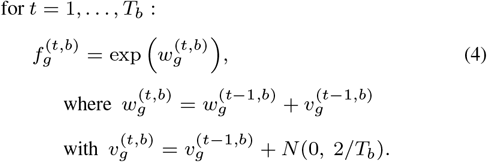

In particular, we initialize

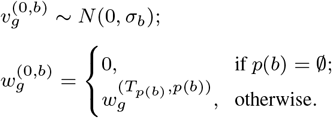

Then, for each branch *b*, we randomly sample several time points to generate the final cell samples, and let the “group” identity of cell sample *c* be *k*(*c*) = (*t, b*).

Finally, we generate the base expression with an adjustment of library size for each gene *g* in cell *c* as

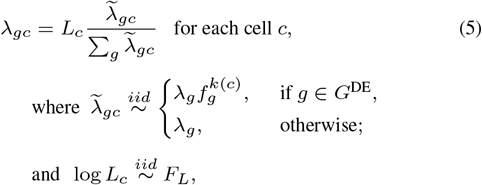

where *k*(*c*) denotes the group identity of cell *c*.

#### Intrinsic variation

##### Marginal distribution

Gene expression in individual cells is an inherently stochastic process (Raj *et al.*, 2006). If the gene regulation is ignored, this process is just a simple two state birth-death process. The steady-state distribution for this stochastic process in most cases turns out to be a Gamma-Poisson, Beta-Poisson, or Poisson, which is justified from the theoretical biochemistry aspect (Grün *et al.*, 2014; Kim and Marioni, 2013), the experimental data sampling aspect (Quinn *et al.*, 2018), and also the common observations from the data. Splat (Zappia *et al.*, 2017) and PROSSTT (Papadopoulos *et al.*, 2019) utilize the negative binomial model in the simulation of marginal gene expression; while SymSim (Zhang *et al.*, 2019a) uses a Beta-Poisson instead; SERGIO (Dibaeinia and Sinha, 2020) simulates the gene expression via solving the series of ordinary differential equation functions following the literature about gene kinetics with regulation (Schaffter *et al.*, 2011).

ESCO adopts the negative binomial model, since it is widely accepted in the literature and enjoys support from biochemistry, experimental data sampling, and empirical observations. Particularly, following Splat (Zappia *et al.*, 2017), we can naturally enforce a mean-variance trend by simulating the Biological Coefficient of Variation (BCV) for each gene. BCV is defined as the square root of the standard deviation divided by the mean, i.e., the square root of the coefficient of dispersion. It has been pointed out (McCarthy *et al.*, 2012) that one should not assume a common dispersion for all the genes, as a gene-specific variation is often detected in RNA-seq case studies. Splat simulates BCV as a weighted sum of a common dispersion and a gene-specific dispersion, such that some information can be shared across genes to benefit the estimation, while preserving the gene-specific variation.

Particularly, we generate the marginal counts 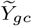 as:

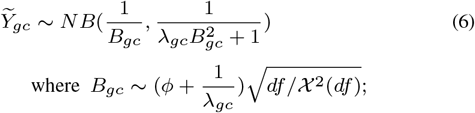

where *ϕ* is the common dispersion parameter, and *df* represents the degree of freedom of the 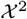, and *NB* represents the Negative Binomial distribution.

##### Co-expression

The gene expression (either the truth or the observed) is not necessarily independent even within cells of the same type, resulting from gene regulation. Characterizing the joint distribution requires solving the steady distribution of multiple correlated stochastic processes, which usually does not have a closed-form solution and requires large computational power (Pratapa *et al.*, 2020; Dibaeinia and Sinha, 2020). Since the marginal distribution of gene expression is understood fairly well, naturally, we think of using the copula to model the gene dependence. This idea is shown to be successful in Inouye *et al.* (2017) to model bulk RNA-seq data.

A copula is defined by a joint cumulative distribution function (CDF), *C*(*u*): [0, 1]^*p*^ → [0, 1] with uniform marginal distributions. One of the most popular copula models is the Gaussian copula, which is defined simply as:

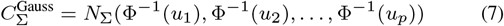

where Φ^−1^ denotes the inverse function of standard normal CDF, and *N*_Σ_ denotes the joint CDF of a multivariate normal random vector with zero means and correlation matrix Σ. Due to the well-known consistency between Σ and the empirical Pearson correlation matrix, the Gaussian copula allows for directly interpretable dependence simulation, and therefore is adopted by ESCO.

Particularly, we generate true counts *Y*_*gc*_ via the following model:

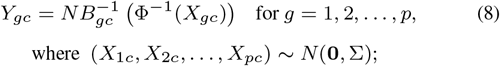

and 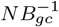 is the quantile function of the Negative Binomial distribution with parameters indexed by cell *c* and gene *g* in equation (6), and Σ is the target correlation matrix.

##### Technical noise

Currently, there are mainly two single cell library preparation protocols: (1) full-length mRNAs profiling without the use of UMIs (e.g., with a standard Smart-Seq protocol); and (2) profiling only the end of the mRNA molecule with the addition of UMIs (e.g., 10x Chromium). The former protocol is usually applied for a small number of cells and with a large number of reads per cell, providing full information on transcript structure. The latter is normally applied for many cells with shallower sequencing, and it is impacted less by amplification and gene length biases. We focus on the UMI-based protocol in this paper because it is usually less biased with greater sparsity.

There currently exist two approaches to simulate the technical noise: one is based on data generating process, and the other is based on data visualization and fitting. As an example of the former, SymSim (Zhang *et al.*, 2019a) uses the empirical approximation of the major steps in the experimental procedures such as mRNA capture, PCR amplification, RNA fragmentation, and sequencing, to directly imitate the technical noise. On the other hand, Splat (Zappia *et al.*, 2017) simulates the technical noise by adopting a zero-inflation model, where the zero-inflation probability relates to the gene expression level in a way that comes from the observed trend in the real data.

There are both pros and cons with regard to these two approaches. The empirical approach facilitates the generation of more realistic noise, but suffers from finding appropriate configuration to match a particular data set (actually, SymSim uses a grid search to do the matching). In contrast, the parametric approach allows a one-step estimation of the parameters from the real data, but can suffer from poor goodness-of-fit due to the mismatch of models. Therefore, ESCO integrates both procedures and gives users the freedom to choose between the two. Particularly, as for the empirical approach from SymSim, one may resort to **Fig. 2 B** and Zhang *et al.* (2019a) for details. While as for the parametric approach from Splat, the observed counts *Z*_*gc*_ from the data is generated via the following

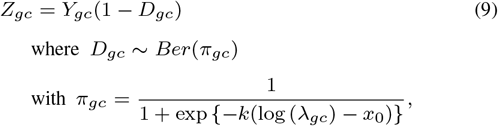

where *π*_*gc*_ denotes the probability of zero-inflation, given the expression mean *λ*_*gc*_, *Ber* denotes the Bernoulli distribution, and *Z*_*gc*_ denotes the final observed counts.

### 2.2 Estimation

ESCO facilitates mimicking any particular data set, consisting of either homogeneous or heterogeneous cell groups, by estimating the hyperparameters from the data. Through learning the parameters in the parametric model, this approach fits data as well as possible (**Fig. 4**), given the limitations of the parametric choice.

**Fig. 4.**
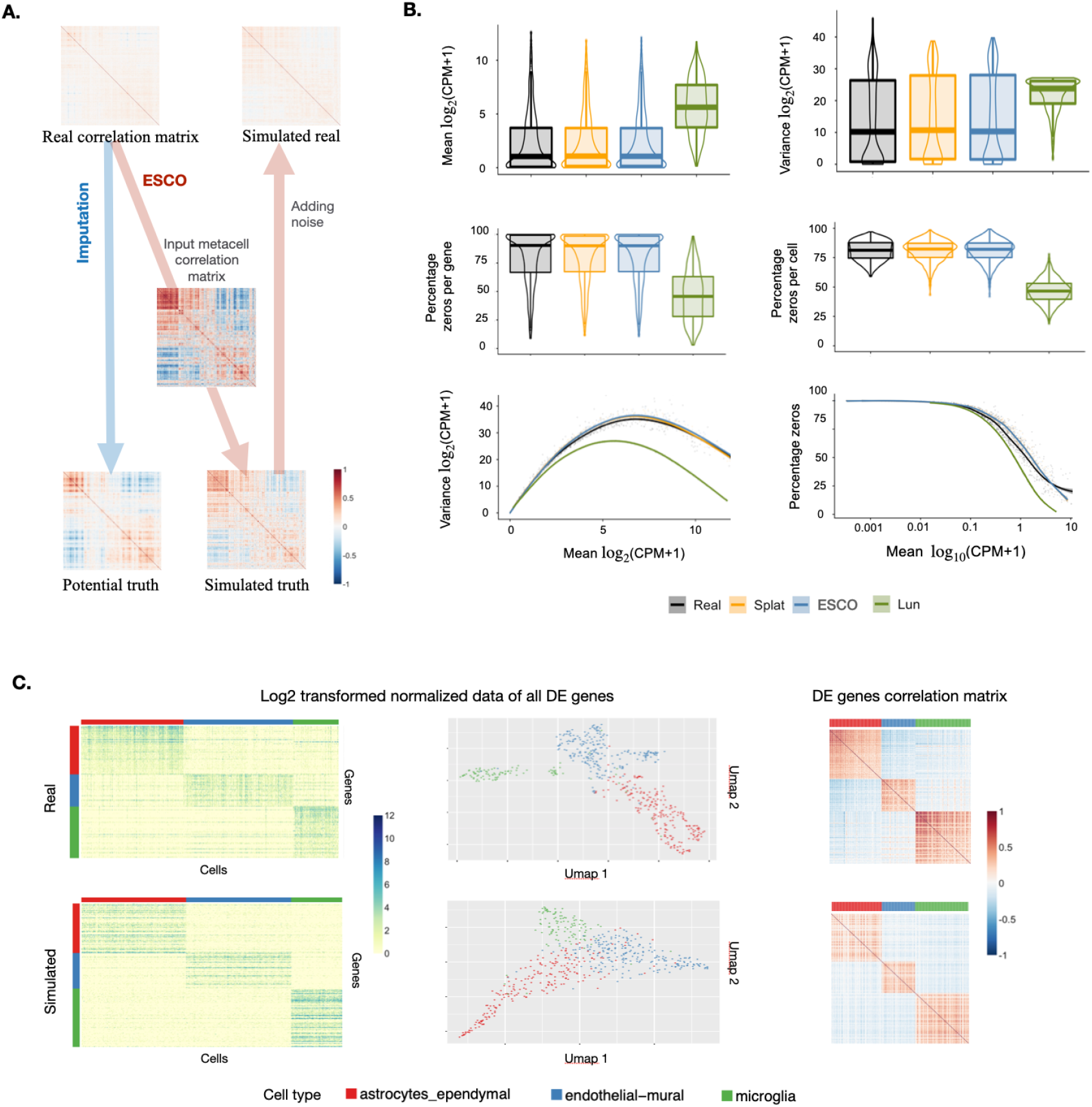
ESCO can learn both the cell heterogeneity and gene co-expression from the data. **A.** The generation process of gene co-expression for one homogeneous cell group from real data using ESCO. Particularly, the example is for 500 randomly selected genes in pyramidal CA1 cell type (911 cells) from Zeisel data. **B.** The comparison of marginal features of real data consist of 500 randomly selected genes in pyramidal CA1 cell type (911 cells) extracted from Zeisel data, and the corresponding simulated data using different simulators. Particularly, Lun (Lun et al., 2016) is one of the earliest scRNA-seq simulators, which has been found to be suboptimal (Zappia et al., 2017). We include it here as a clear contrast with the state-of-art methods. **C.** The comparison of real data consist of 4000 most differential expressed genes in three cell types (astrocytes_ependymal, endothelial_mural, microglia) of 526 cells in total extracted from Zeisel data, and the corresponding simulated data using ESCO. While the UMAP depiction differs somewhat, the expression and co-expression patterns match closely.

Next, we elaborate on our specific estimation strategies. Recall that ESCO takes a hierarchical modeling approach, paired with a copula. As such, an empirical Bayesian approach to parameter estimation would be appropriate. However, it is usually infeasible to compute the solution. Therefore, we follow Splat and estimate the parameters in each layer separately. Particularly, we assume the data are already normalized (i.e., no batch effect arises due to technical reason) and have disjoint marker gene sets across cell types, and consider the three estimation tasks in the following.

#### Estimating the heterogeneity

We have introduced three types of heterogeneity of gene expression (discrete, tree, and trajectory), but we only present an estimation procedure for the discrete one here, leaving the more complex structure of the other two models to future work. Following our modeling of the discrete heterogeneous cell groups, we first split all the genes to DE and non-DE genes based on their AUC scores in cell group prediction using SC3 (Kiselev *et al.*, 2017), provided that we already have the true cell group annotation. Particularly, we use 0.7 as our cutting threshold of the AUC score, i.e., classifying the genes with AUC score no less than 0.7 as DE genes and the others as non-DE genes.

We then use the DE genes to estimate the DE factors. Particularly, we divide those DE genes into marker genes for each cell group based on their classification result from SC3 (Kiselev *et al.*, 2017). We assume that the mean distribution of marker genes in their marked cell group follows the same distribution in the other cell group and a DE factor that follows LogNormal distribution indexed by the cell type. Therefore, we estimate the DE factor for marker genes of cell group *k* via fitting a LogNormal distribution on the ratio of their sample mean within cell group *k* and those outside cell group *k*.

#### Estimating the intrinsic variation

##### Marginal

As for estimating the parameters related to marginal intrinsic variation, we follow the technique used in Splat (Zappia *et al.*, 2017), with a few refinements. We allow non-parametric fitting of the library size distribution and base mean distribution, which can be done quickly by computing the empirical CDF and also later on sampled from using Metropolis-Hastings sampling due to the univariate nature. One may refer to Zappia *et al.* (2017) for further details about the estimation procedure for other marginal parameters included in the algorithm, such as BCV and outlier.

##### Copula

To circumvent challenges due to technical noise and sparse counts, we cluster similar cells and form metacells (Baran *et al.*, 2019) and then estimate Σ for equation (8). As an integrated version of the original real data, the size of metacells must be carefully selected so that the technical variation can be reduced, while some biological variation can be preserved. We refer the reader to the source paper of MetaCell (Baran *et al.*, 2019) for further details.

A more statistically convincing approach would be the non-parametric estimation procedure called SKEPTIC (Liu *et al.*, 2012), which is built for a continuous marginal paired with a Gaussian copula. However, SKEPTIC is derived assuming a continuous marginal without additional noise. In our case, the data are discrete, and the underlying truth is severely masked by the additional zeros, so we find it challenging to recover signals from real data. Therefore, we did not consider this direction, though careful adjustment of the estimation procedure and corresponding consistency under the discrete marginals masked by false zeros is worth attention in future work.

#### Estimating the technical noise

ESCO also allows estimation of the median zero-inflation and shape parameters in equation (9). Though Splat already includes the corresponding estimation via fitting a logistic regression between the log-transformed gene mean and their observed zeros proportions, it is biased towards inflating the probability of excess zeros, as can be understood via the following reasoning:

Given a real scRNA-seq data set 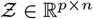, where each element *Z*_*gc*_ is the observed count of the expression of gene *g* in cell *c*, let

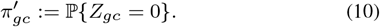

Splat estimates 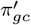 via fitting a logistic function to model the relationship between the log means of the normalized counts and the proportion of cell samples that are zero for each gene. Then Splat plugs the estimation 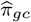 in place of *π*_*gc*_ in equation (9) to simulate 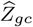,

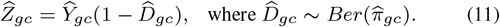

and 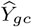 is the imitation of the true counts *Y*_*gc*_ for gene *g* in cell *c* simulated in the previous steps.

Assuming the estimation of 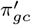 is accurate and the simulated true counts 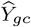 well mimics the real truth *Y*_*gc*_, then this approach would cause more sparsity than expected, since the proportion of zeros in the simulated observation will be

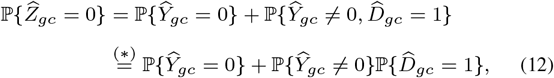

where (*) is true since 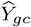 and 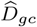 are independent once condition on *λ*_*gc*_. Therefore,

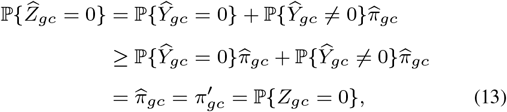

From the above calculation, one simple correction for this bias uses:

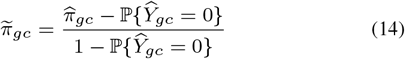

as the plug-in for equation (9). Particularly, ESCO approximates 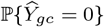 using the CDF of Poisson with mean *λ*_*gc*_ at zero.

## 3 Results

Recall that a particularly prominent aspect of noise that complicates scRNA-seq data analysis is sparsity due to low capture and sequencing efficiency in the data collecting process. Excess sparsity has been shown to corrupt the analysis of scRNA-seq data in many ways (e.g., cell clustering, trajectory inference, DE gene detection, etc.). Imputation methods can generally help according to several benchmarking efforts (Zhang and Zhang, 2018; Andrews and Hemberg, 2018). However, the influence of sparsity on gene co-expression, particularly within the homogeneous cell group, has been overlooked by many. ESCO provides an easy way to fill in the gap, as it allows for the generation of flexible gene co-expression as a ground truth. In the following we present a systematic evaluation of the performance of imputation methods on the recovery of gene co-expression using ESCO.

### 3.1 Sparsity attenuates the gene co-expression

First, we show that sparsity indeed impedes the recovery of gene co-expression in scRNA-seq data. Highly expressed genes are much less likely to suffer from technical noise, as they have sufficient replicates to be detected in the data collecting process, in contrast to relatively lowly expressed genes. To illustrate this point we contrast gene co-expression for marker genes in scRNA-seq data (Velmeshev *et al.*, 2019) to bulk RNA-seq data (Parikshak *et al.*, 2016). Genes are classified as high or mid, based on their expression values. In scRNA-seq data, the mid-genes demonstrate substantially less correlation when compared to the high-genes (**Fig. 5A** top panel). But in the bulk RNA-seq data, mid and high-genes demonstrate equivalent levels of correlation (**Fig. 5A** bottom panel). Because we expect little, if any, impact of technical noise in bulk data, and similar levels of correlation for marker genes in these two data sources, this investigation suggests that sparsity attenuates measured correlation of gene expression in scRNA-seq data. Thus we look to imputation for improved performance.

**Fig. 5.**
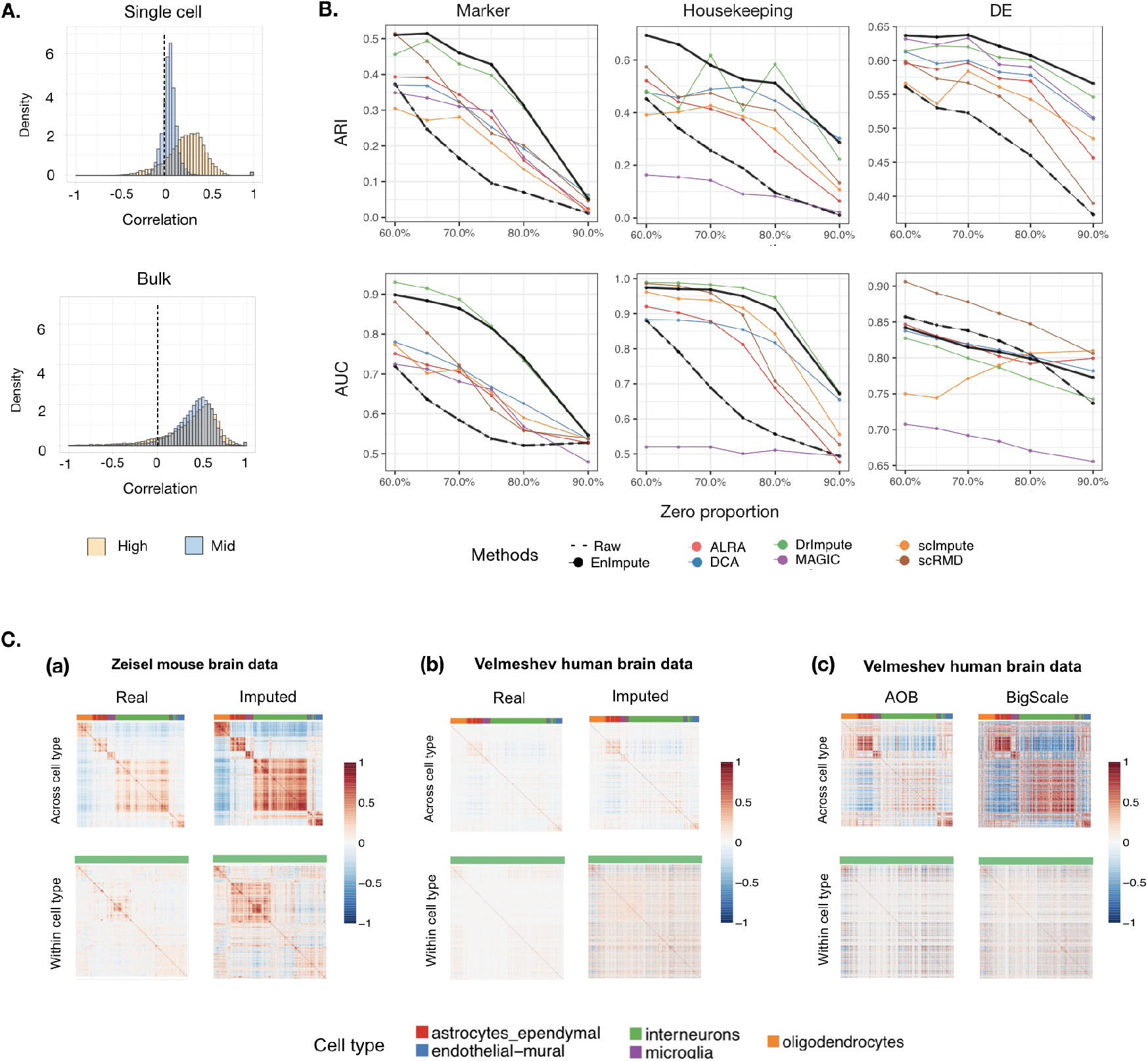
Application of ESCO in benchmarking imputation for gene co-expression recovery. **A.** Evidence that sparsity attenuates gene co-expression. The top panel depicts the histogram of Pearson’s correlations for the 1000 highest expressed (≈0-10% quantile) genes and 1000 moderately expressed genes (≈60-70% quantile) in Velmeshev scRNA-seq data. The bottom figure depicts the histogram of Pearson’s correlations for the same genes as in the top panel, but using the corresponding bulk data. **B.** The performance of different imputation methods on recovering the gene co-expression. We simulate 1000 genes and 200 cells for three cell groups, using the parameters estimated from the Zeisel data, and aggregate the results from 10 replicates. The corresponding ARI score and AUC score (represented by each row) of each imputation method versus different sparsity levels (represented by zero proportion) on different types of gene co-expression (represented by each column, respectively, as marker genes, housekeeping genes, DE genes) are plotted. **C.** Verification of the findings of imputation using real data. **(a)** The correlation matrix of marker genes before and after imputation of Zeisel data, across cell types (five in total) and within one cell type (interneurons). **(b)** The correlation matrix of marker genes before and after the imputation of the Velmeshev data. **(c)** The correlation matrix of marker genes of the Velmeshev data after AOB and BigScale aggregation.

### 3.2 Imputation can help recover GCN with moderate sparsity

Working with Zeisel data (Zeisel *et al.*, 2015), we consider a subset of data consisting of the 4000 most differentially expressed genes and 526 cells from three cell types (astrocytes_ependymal, endothelial-mural, microglia) that have distinct marker genes. We simulate data from 1000 genes and 200 cells with hyperparameters estimated from the real data, while manually changing the sparsity level such that the zero proportion ranges from 60% to 90% (the real data has ≈ 43% zeros), and aggregate the results over replicates. We evaluate ARI and AUC score for each imputation method under a range of sparsity levels (i.e., zero proportion) for various gene sets of interest (**Fig. 5B**). Specifically, we choose the number of clusters in ARI, calculated via a grid search over a range of clusters numbers (2-9 in our case), with the highest score. Additionally, we calculate AUC by assigning gene pairs as connected or un-connected based on the co-expression significance in permutation testing of the simulated truth. We then assess the prediction accuracy (AUC scores) of connections for each imputation methods using their estimated co-expression. All the results are averaged over 10 replicates. We observe that:

1. generally, when facing moderate sparsity, all the imputation methods beat the un-imputed raw data, depicted by the bold dashed black line;
2. the performance of imputation gets worse with excessive sparsity;
3. as for a comparison among different methods, there is no universal winner for all settings, but the ensemble method, depicted by the bold black line, provides the best or close to the best performance across all settings we considered.

In the following section, we aim to verify our findings of imputation using real scRNA-seq data. It is conjectured that the co-expression of marker genes in the mouse brain will be similar to that of the human brain. Therefore, we expect the recovered gene correlation from a data set measuring mouse brain will follow a similar pattern to those from the data set measuring the human brain. Particularly, we use Zeisel data (Zeisel *et al.*, 2015) for the mouse brain and Velmeshev data (Velmeshev *et al.*, 2019) for the human brain. The Zeisel data have deeper sequencing for single cells and consequently are less noisy, with less sparsity, compared with the Velmeshev data, which have a much greater number of nuclei sampled, each with fewer reads. Therefore, we can see the influence of the sparsity level on gene co-expression by directly comparing these two data sets. We select five common cell types in both data sets and use the Zeisel data as the benchmark. We evaluate the correlation matrix of marker genes before and after imputation of Zeisel data, across cell types and within one cell type (i.e., interneurons). **Fig. 5C(a)** plots both the gene co-expression across heterogeneous cell groups and gene co-expression within homogeneous cell groups before and after imputation with EnImpute method (Zhang *et al.*, 2019b) using Zeisel data, while **Fig. 5C(b)** plots the same results but using the Velmeshev data. We can see that for the Zeisel data (moderate level of sparsity), imputation enhances the gene co-expression pattern both within homogeneous and across heterogeneous cell groups. In contrast, for the Velmeshev data (excessive sparsity), imputation fails to help much to recover the gene co-expression across heterogeneous cell groups pattern, while failing utterly for gene co-expression within homogeneous cell groups, which is expected, as it is a harder task. This investigation supports some of our findings of imputation, i.e., imputation can generally help, but may fail as sparsity levels increase to a very high level.

### 3.3 Data aggregating is a better way to recover GCN with excessive sparsity

Despite the excessive sparsity in the Velmeshev data, these data have the advantage of abundant numbers of cells, which inspired us to explore another approach for recovering gene co-expression: data aggregation that utilizes the abundance of measured cells. We introduce two methods below, a simple heuristic (AOB) and a complex algorithm (BigSCale).

#### Averaging over cell bags

If one has successfully assigned the cell type labels, one may be able to use the simple procedure of averaging gene expression over random splits within cell types, and then compute the gene co-expression based on those averaged values (Polioudakis *et al.*, 2019). We will refer to this procedure as AOB (**A**veraging **O**ver **B**ags). The only tuning parameter here is the bag size, which should be chosen carefully so that we can mitigate the influence of sparsity and other noise, while still maintaining some variability among samples.

#### Pre-clustering and transforming the expression value

More recently, a method called BigSCale (Iacono *et al.*, 2019) was developed for the problem of recovering GCN in a similar, but more complex way. This algorithm first clusters cells sharing highly similar transcriptomes together, and then treats them as biological replicates to evaluate the noise and an indirect measure of correlation. This method works well when there is a sufficiently large number of cells for meaningful cell clusters to form, but it is pretty computationally challenging.

We find both methods work well in recovering gene co-expression across heterogeneous cell groups (**Fig. 5C(c)**), though neither successfully recover gene co-expression within homogeneous cell groups. Future efforts are needed to recover these subtle signals.

## 4 Discussion

In this paper, we propose a new scRNA-seq simulator, ESCO, which borrows the good features of the current state of art simulators in an ensemble, while for the first time, allowing both interpretable and controllable gene co-expression generation. Specifically, ESCO allows realistic simulation of various cell group structure, ranging from simple homogeneous cell groups to tree-structured discrete cell groups to continuously changing cell trajectories, together with gene co-expression. ESCO outperforms other methods as it preserves the highlights of all the other existing simulators in one R package, including the hierarchical semi-parametric modeling of homogeneous groups from Splat, the tree-structure generation from SymSim, and the trajectory generation from PROSSTT, all while interjecting gene-gene interactions. Specifically, ESCO allows the flexible generation of both gene co-expression across heterogeneous cell groups arising from a cell group structure and gene co-expression within homogeneous cell groups arising from gene-gene interaction in one functional cell group, which have been overlooked and underdeveloped in other methods.

There is still much room for future work in this area. The efficient estimation of the hyperparameters from the real data in the tree-structured cell group and continuous cell trajectories scenario still needs improvement. Currently, most simulators rely on a grid search of parameters to find parameters that fit a particular data, but these parameter choices do not extend to new settings, and it is extremely challenging to simulate data similar to new data sets. The ability to simulate realistic batch effects in various settings is also not satisfactory in the current methods. ESCO, which mimics Splat in this regard, shares this shortcoming. A careful, deep-dive to produce realistic batch effects is needed.

## Footnotes

1 SymSim generates the gene expression for gene *g* in cell *c* via a random product model, that is expression *Y*_*gc*_ = *λ*_*g*_*τ*_*c*_, where 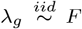, and 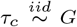. Once conditioning on the cell confounder *τ*_*c*_, the correlation between expression of genes *g*_1_ and *g*_2_ disappears.

